# Shaping coral traits: plasticity more than filtering

**DOI:** 10.1101/2021.08.19.456946

**Authors:** Viviana Brambilla, Miguel Barbosa, Inga Dehnert, Joshua Madin, Davide Maggioni, Clare Peddie, Maria Dornelas

## Abstract

The structure of ecosystems is usually determined by the shape of the organisms that build it, commonly known as ecosystem engineers. Understanding to what extent plasticity and environmental filtering determine variation in ecosystem engineer physical structure is necessary to predict how ecosystem structure may change. Here, we explored coral survival and the plasticity of morphological traits that are critical for habitat provision in coral reefs. We conducted a reciprocal clonal transplant experiment in which branching corals from the genus *Porites* and *Acropora* were moved to and from a deep and a shallow site within a lagoon in the Maldives. Survival and trait analyses showed that transplant destination consistently induced the strongest changes, particularly among *Acropora* spp. The origin of the corals only marginally affected some of the traits. We also detected variation in the way individuals from the same species and site differentiate their shape, showing that traits linked to habitat provision are phenotypically plastic. The results suggest coral phenotypic plasticity plays a stronger role than environmental filtering, in determining zonation of coral morphologies, and consequently the habitats they provide for other taxa.

## Introduction

Organismal morphology can vary across environmental gradients. This variation can be due to specific morphologies suiting particular conditions (environmental filtering) or organisms developing different morphologies in response to different conditions (plasticity) (1). Physical ecosystem engineers create the habitats available within an ecosystem, and for this reason their morphological variation has strong implications for ecosystem function (2). Here we investigate the relative role of environmental filtering and plasticity on coral morphology, focusing on continuous morphological traits that are linked to habitat provision on coral reefs. Whether the coral morphologies we observe result from filtering or plasticity will give us an idea of the extent to which reefs rely on coral plasticity for functioning, and how much habitat variability is due to organismal properties rather than broader environmental constraints (3).

Ecosystem engineers are organisms whose presence or activity alters their physical surroundings or changes flow of resources within the local populations (2). Plants, for example, modify habitats within a forest with their structures (4) and act as chemical engineers by altering soil nutrients with their root system for the whole soil ecosystem (5). In the marine realm, some organisms act as physical ecosystem engineers through the accumulation of bio-constructed carbonate calcium structures. For example, some algae deposit calcium carbonate formations, some bivalves build shell beds, and corals have hard aragonite skeletons that shape reef substratum (2). Structures built by ecosystem engineers give a third dimension to otherwise almost flat surfaces and affect local environmental conditions, in turn increasing microhabitat diversity. Understanding what determines variation of ecosystem engineers morphology is then important to understand ecosystem dynamics and evolution (6), and may help understand the extent to which organisms can change the environment they experience (7).

Plasticity and environmental filtering are important drivers of engineers’ morphological variation. Environmental filtering is a form of natural selection, whereby environmental conditions prevent some phenotypes from persisting in some places (8). If applied to engineering species, it means that the environment favours some type of habitat constructors. Phenotypic plasticity, however, may override the constraints of environmental filtering, by allowing the same genotype to develop phenotypes that are better suited to different environmental conditions (9). When environmental conditions change, ecosystem engineers can perish (filtering) or adapt to the new condition through plasticity. Phenotypic plasticity and filtering can therefore simultaneously contribute to variation in prevalent morphologies in an assemblage of engineers. Therefore, local habitat conditions result from both a bottom-up (habitat construction via ecosystem engineers) and top-down (natural selection and environmental filtering) processes. If effects of plasticity are greater than those of environmental filtering, then organisms play a primary role in determining ecosystem functioning and can play an active role in countering effects of natural selection (3). To test this hypothesis coral reefs are optimal, as reef habitats are shaped by the hard corals inhabiting them.

Scleractinian corals are primary ecosystem engineers on most coral reefs. Hard corals accumulate calcium carbonate through skeleton accretion, which is the basis of the reef matrix. Since most corals rely on light for energetic intake through photosynthesis, and photosynthesis is tightly linked to calcification, morphological differences are particularly evident along light gradients (10–14). In nature and for a number of species, flattened and horizontally developed morphology tend to develop in deep environments where there is less light (15), possibly to minimize tissue that has to be sustained for any given light flux (16,17). On the one hand, reef zonation is very evident along environmental gradients such as depth or wave exposure gradients (18). On the other, corals can respond to environmental factors by changing their morphologies in ways that are often found convenient for their fitness. For example, they can maximize physiological efficiency (17) or particle capture (19) and be present under different environments but developing different structures. Yet, how much of this variation in shape is driven by environmental filtering vs. plasticity, and how this balance differs across species, remain poorly understood.

Reciprocal transplant experiments allow following transplants’ phenotypic response within generational time and drawing phenotypic reactions of exposure to different environmental gradients (*i*.*e*, reaction norms, 12). However, to disentangle the drivers of variation in coral morphologies, we need to quantify morphologies themselves. Scleractinian corals’ morphologies are traditionally divided into discrete growth forms based on qualities of their shapes. For example, the presence and structural organization of branches, the tendency to “encrust” substrata, or the build of bulging morphologies. (20,21). Using categories does not allow capturing intra-category variation in morphologies, nor how different morphologies translate into physiologically and ecologically relevant variation (22,23). To capture these differences, the use of quantitative continuous variables coupled with defined ecosystem functions allows describing different coral morphologies along continuous axes of variation and across growth forms. For example, colony compactness promotes reef stability, surface complexity promotes microhabitat diversity and recruitment facilitation and top heaviness provide large fish refuge (23). Each one of these variables can be measured continuously by focusing on the 3D geometric aspects that define that property (22). In this study, we quantify the drivers of variation across multiple axes of coral morphology, including those along these niche constructing traits.

So far, categorical, discrete or corallite morphological features of corals have been showed to be affected by filtering, plasticity and/or adaptability under a wide range of gradients (12). Of all, light is the factor that most consistently correlates with morphological features (15,24). For example, corallite architecture of *Goniastrea pectinata* changes with light exposure so that the induced calyx morphology under higher exposure is efficient at shading, while under lower exposure is efficient at maximizing light capture. As for most natural populations, genotype by environment interaction is often detected when studying coral plasticity studies (12,15,24–26). Yet, quantification of variation in plastic responses at individual, population, and species level, and how much potential there is for environmental filtering to act on coral evolutionary pattern, is rarely tackled. To gain a stronger understanding of the role of plasticity and environmental filtering in coral evolution, comparisons of quantitative traits that continuously characterize colony shapes and functions within and across taxa with phylogenetic dependencies are needed.

This study aimed to partition the variation in coral morphology driven by environmental filtering and plasticity among differently related species. To achieve this aim, we used a reciprocal transplant experiment using taxa of the same broad category of morph type (branching) but belonging to two different genera. We tested if: i) coral colonies from different origins differ in their change in morphology (evidence of environmental filtering); ii) different environmental treatments consistently induce different structural morphologies (evidence of plasticity); iii) differences in the direction of the transplantation, this is pairs of origin and destination site, induced different morphologies (evidence of local adaptability); and iv) genus, species or genotype affected change in coral morphologies and the extent of their effects (evidence of evolutionary constrain and genotype x environment interaction).

## Methods

### Site, species and transplant

A coral reciprocal transplant experiment was set up in the South-East lagoon of Maghoodoo Island (3°04′N, 72°57′E, Republic of Maldives) from January 2017 to May 2018. Transplants were made between a shallow (S) high-light site at 5-6 m depth and a deep (D) low-light site at 16-18m depth. Five replicate racks for coral samples were built and fixed to the reef at each site.

Coral samples were collected at the two sites from four coral species with different branching morphologies. The species were *Acropora divaricata* (arborescent/corymbose), *Acropora muricata* (arborescent), *Porites rus* (encrusting/digitate), and *Poritis cylindrica* (digitate) (Figure 1). Species were chosen from two genera to test if plasticity is similar across evolutionary lineages. All species were found at both sites, were easy to identify, and had relatively high growth rate (27).

**Figure 1 -.**
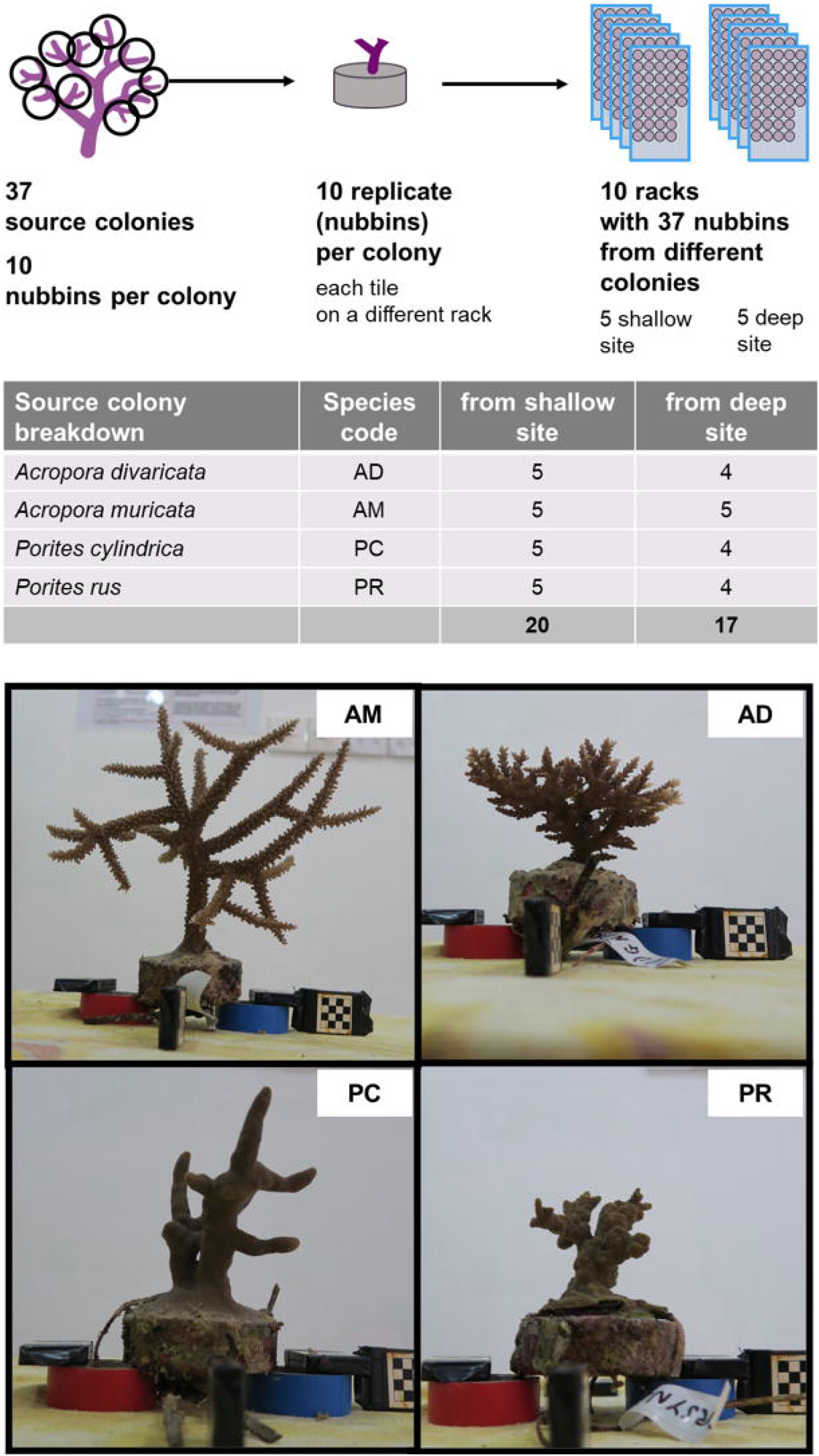
Reciprocal transplant experiment schematic and details. On the top, the experimental design followed in January 2017 is shown, with details of the 37 source colonies’ origin (table). Pictures on the bottom show specimens for the species used in the experiment. Pictures were taken at the end of the experiment (May 2018).

For each species, 4-5 source colonies were selected at each of the shallow and deep sites, for a total of 37 colonies (Figure 1). Colonies were a minimum distance of 10 m apart to avoid collection of colonies with the same genotype (clones) (28). Colonies were transported to the lab and kept in aerated tanks. Ten replicate nubbins were pruned from each colony: five to be returned to the source site (transplanted within site as a control), and five to go to the other site (transplanted between sites). Nubbins were cemented to a concrete disk tile (7 × 2.5 cm) with reef cement (NYOS © reef cement). The maximum basal diameter (D, cm), minimum basal diameter (d, cm) and length (L, cm) of each nubbin was measured with callipers. Wet weight (*i*.*e*., the weight of the nubbin on the tile as it was taken out of the tank, W, in grams) was measured at the closest 0.5 g. Volume (V, mL) was measured with the water displacement method with graduated cylinders (29). Photographs of the nubbins were taken from above at a fixed distance with a scale bar. Each source colony was processed and transplanted back to the reef within 24 hours. All the samples (n=370) were collected, processed, and transplanted in six days. One year and 4 months later (May 2018), all the racks were brought to the lab, where the status of each nubbin (alive or dead) was recorded, and the same set of measurements and photographs were repeated.

### Environmental data collection

Environmental data—sediment, temperature, water flow and light—were measured at each site. Sediment traps were built and deployed as per English et al. (1997) in April 2017. Three traps per rack were deployed for 10 days (9th to 19th of April) and checked to make sure there was no overgrowing on their top.

Flow was measured with the gypsum balls method, which is based on the fact that gypsum dissolving in water is directly proportional to flow velocity (30). The gypsum balls were weighted before being deployed in the field for a 24-h period (+/-20 min). At each site (*i*.*e*., shallow and deep), we allocated three gypsum ball replicates, resulting in six replicates per depth. After retrieval, each ball was left to dry for two days in a low-humidity room at 25°C and then weighted again. A proxy for flow states was obtained as the weight lost during deployment.

We also allocated HOBO temperature and light dataloggers (Onset) to each rack. These were collected at the end of the experiment. At least two loggers for depth were deployed at all times and they were replaced three times along the experiment. Since loggers became covered by fouling organisms, only the first month of each deployment was used to measure light treatment per site. Since temperature loggers tend to accumulate heat when exposed to light, only the last month of each deployment (*i*.*e*., when the logger was covered with a biofilm protecting the logger from solar radiation) was used to estimate temperature.

### Photograph processing and morphological traits

Nubbin photographs were processed using the image analysis software ImageJ (31). A graphics tablet (medium Intuos, Wacom) was used to draw the outline of the planar projection of the nubbins on the horizontal plane. Using the scale bar present in the picture as a reference, the contours were then saved as XY coordinates to calculate planar area (PA, in cm^2^) and perimeter (p, in cm) with the ‘pracma’ R package (32). Furthermore, 2D metrics to reflect three aspects of coral morphology (top-heaviness, surface complexity and shape compactness) were calculated. To measure top-heaviness, the log-ratio between the radius of the best fit circle to the outline of the nubbin and the mean basal diameter was computed (T, cm). To measure shape compactness (C, non-dimensional), the circularity formula was used:

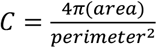

To measure shape complexity, two non-dimensional indexes were used. Rugosity index (R), measured as the ratio between the outline perimeter and the perimeter of a circle with the same area as the outline, and Fractal dimension (F), measured with the ‘fractaldim’ package (33) and specifying ‘box counting’ as method.

To visualize how traits changed collectively during the experiment, a principal coordinate analysis (PCA) considering all the observed combination of traits (*i*.*e*., pooling nubbins from start and end of the experiment) was performed with the ‘prcomp’ function. Traits were previously log transformed and the multidimensional space was used to inspect how different groups of nubbins were occupying different portions of the morphospace.

### Analysis

To analyse probability of survival, nubbin status (alive - 1 or dead- 0) was used as response variable and a binomial generalized linear model with logit link function was used. To measure change in morphological variables, log-ratios (lr) of the variable between the end and the start of the experiments were used. To measure change in the morphospace, vectors from the initial to the final positions of each nubbin in the space defined by the first two principal components were analysed. Specifically, distance (deltaPCA) defined as the module of the vector, and direction (dirPCA), defined as angle of inclination of the vector, were quantified.

Generalized and linear mixed models were used to examine the effect of coral colony origin, transplant destination and genus on each of survival, maximum and minimum diameter, nubbin length, volume, weight, planar area, compactness, surface rugosity, fractal dimension, top-heaviness, delta PCA and direction PCA. Model selection based on AIC was used as described in Zuur *et al*. (2009). Effects of coral Origin, transplant Destination and Genus were interpreted as effects of environmental filtering, phenotypic plasticity, and taxonomic group, respectively. Genus was used as taxonomic grouping factor because each pair of species is visibly more similar morphologically within Genus groups. Full models included a triple interaction of the three factors of interest because: (i) taxa were expected to be affected differently by the environment, and (ii) we were interested in detecting the interaction between Origin and Destination as signal of local adaptability. Species, genotype and rack were all initially included as random effects. To select the appropriate combination of random effects, full models comprising every possible combination of random variables were fitted with restricted maximum likelihood and the random effect structure corresponding to the full model with the lowest AIC was selected (34). For each response variable (survival, maximum and minimum diameter, nubbin length, volume, weight, planar area, compactness, surface rugosity, fractal dimension, top-heaviness, deltaPCA and dirPCA) and maintaining the selected random effect structure, 18 models comprising every possible combination of fixed effects and interactions were fitted and selected based on the lower AIC. Model assumptions were evaluated graphically (34).

To validate results, we also report the significance of each factor of each full model, with significance levels adjusted to account for multiple comparisons (Bonferroni correction, n = 8). Statistics were performed with R version 3.3.2 (35) and the ‘lme4’ package (36).

## Results

### Environment

There was greater daily variation in temperature and light at the shallow site than at the deep site. While mean temperature differences between depth were not significant (n = 25, df = 23, F value = 0.524, p = 0.477), light values (lux) were higher at the shallow site (n = 25, df = 23, F value = 19.56, p < 0.001), where light levels were 2.2 times higher than at the deep site.

Even though sedimentation rates were not significant between sites (n = 10, df = 8, F value = 2.773, p = 0.134), the shallow site mean (0.031 g/d, sd = 0.017 g/d) was double that of the deep site (mean = 0.015 g/d, sd = 0.013 g/d). Gypsum dissolution varied significantly between depths (n = 12, df = 10, F value = 16.24, p = 0.002), showing that the shallow site had higher water flow. However, the amount of gypsum dissolved during the deployment was smaller than the minimum recorded by other studies during the calibration procedure (30), suggesting that overall, the conditions were of very low water motion at both sites (close to 0 cm/s). This was expected since the lagoon is protected by a high outer reef and as result, only the tidal variation is the main source of water motion, affecting the shallow site more than the deep one.

### Survival

Overall, approximately 60% of the transplanted nubbins survived (Figure 2a). We failed to detect an effect of site of origin on probability of survival, however, there was a significant interaction between genus and destination (Figure 2b, c). Transplanted nubbins were less likely to die if moved to the deep site. This difference was more prominent for the genus *Acropora* spp. (Figure 2c).

**Figure 1 -.**
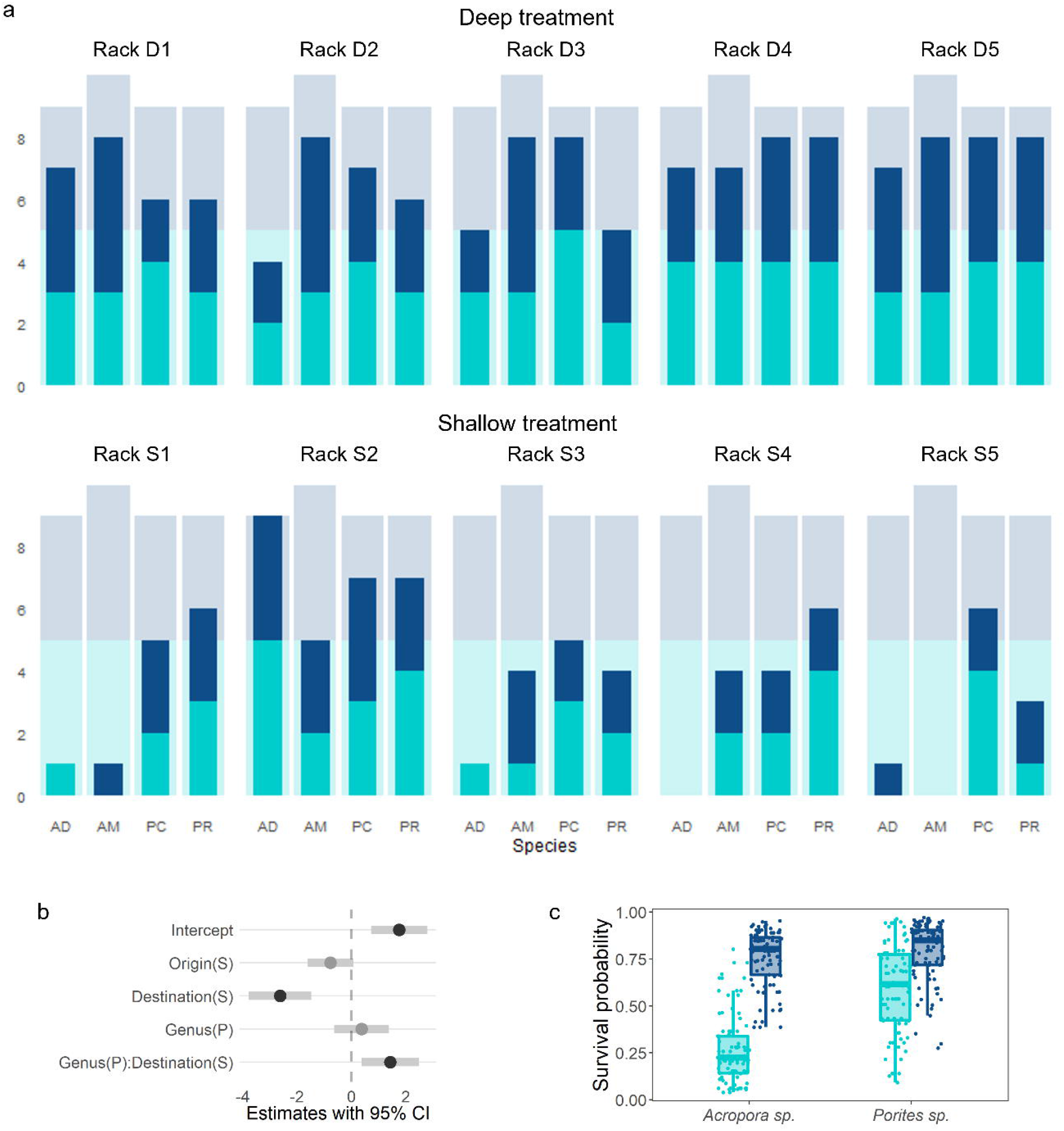
Nubbin survival and model results. *a)* Counts of survived transplant divided by species (AD = *Acropora divaricata*, AM = *Acropora muricata*, PC = *Porites cylindrica*, PR = *Porites rus*) and color-coded by site of Origin (light blue = shallow site, dark blue = deep site). On the top, survivorship among racks deployed at the deep sites. On the bottom, survivorship among racks deployed at the shallow sites. Bulk-colour bars represent counts of the nubbins survived. Shaded in the background, counts of the nubbins originally transplanted on each rack. Differences between the bulk and the shaded columns represent nubbin deaths. *b)* Effect size estimates for fixed factors and interactions for the selected model. Dots represent the estimated effects of each fixed variable and bars represent 95% confidence intervals. An effect is detected (black dots) when bars do not overlap 0, the vertical dotted line. *c)* Survival probability distributions across Genera and Destination (light blue = shallow site, dark blue = deep site).

### Trait changes, reaction norms and morphospace

Changes in morphological traits during the experiment among species and treatments are shown in Figures SM2 and SM3. The first two principal coordinate axes of the continuous morphospace built from the morphological traits explained 84.5% of the variation in nubbin shape (Figure 3a, SM5 and SM6). Nubbins at the beginning and at the end of the experiment occupied different areas of the morphospace (Figure 3a). The initial area occupied was smaller than the final one, as fragments transplanted at the beginning of the experiment were as similar as possible across all experimental nubbins. Differences where more suble when comparing origin and destination groups (Figure SM7).

**Figure 3 -.**
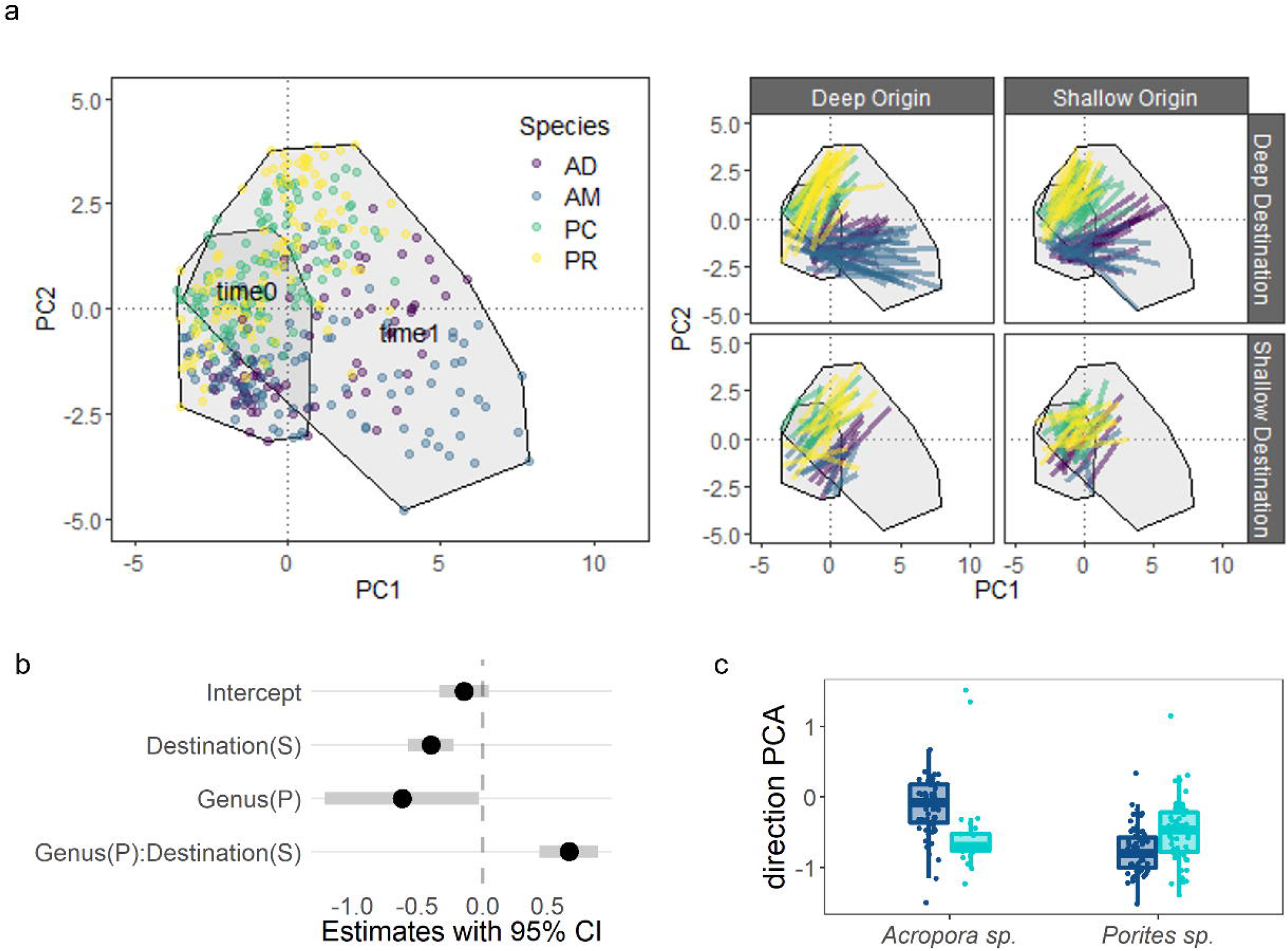
Morphological traits space plot. A) Dots are color-coded by species (AD = *Acropora divaricata*, AM = *Acropora muricata*, PC = *Porites cylindrica*, PR = *Porites rus*). In the space, the 2 polygons represent the space occupied by the nubbins at the beginning of the experiment (time0) and at the end (time1). b) Segments represent the distance travelled by each transplant during the experiment, color-coded by species. The data is shown divided by Origin (columns) and Destination (rows). Trait loadings are shown in Figure SM3. c) Effect size estimates for fixed factors and interactions for the selected model for vector direction. Dots represent the estimated effects of each fixed variable and bars represent 95% confidence intervals. An effect is detected (black dots) when bars do not overlap 0, the vertical dotted line. c) Direction distributions across Genera and Destination (light blue = shallow site, dark blue = deep site).

### Models results

All but two models included a significant interaction between Destination and Genus (Figure 4, Table SM1 and SM2). This indicates that the two taxonomic groups respond differently to the environment they are exposed to. For all traits except basal diameter, trait changes between sites were greater among individuals of the genus *Acropora*. A small effect of Origin was found in changes in nubbin length, planar area and distance travelled in the morphospace (deltaPCA). However, the interaction between Origin and Destination was not significant, suggesting that differences in traits did not depend on the transplant direction (Figure 4, Figure SM8 and Table SM2).

**Figure 4 -.**
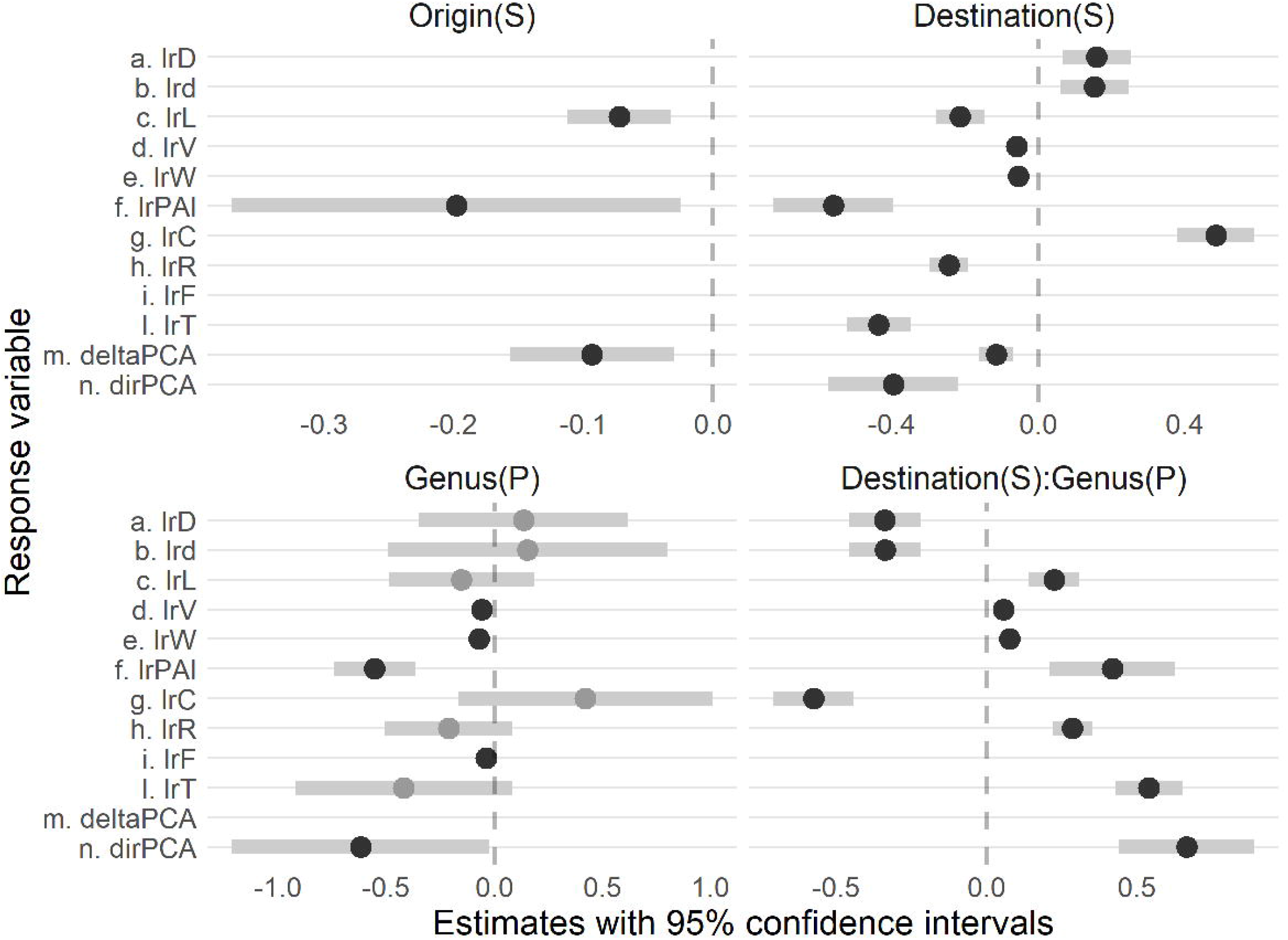
Effect size estimates for fixed factors and interaction terms included in the selected models. Dots represent the estimated effect size of each fixed term in each model and grey bars represent the 95% confidence intervals. Effects are significant (black dots) when bars do not overlap 0 (vertical dotted line in the plots). Notice that the scale varies across panels. P = *Porites* sp.; S = Shallow site.

Detailed results from the models fitted for each response variables are available in the Supplementary Material (see “Models selected for each individual trait”). When looking at the change in the position of the nubbins in morphospace (Figure 3a), each species tended to occupy different region of the space. The direction travelled by each individual (dirPCA) depended on the interaction between environment of exposure and Genus, and was not affected by the Origin of the transplant. Differences were particularly visible in the *Acropora* spp., when comparing Destination sites (higher distance travelled in the deep site), while differences between the *Porites* spp. were smaller (Figure 3b and 3c). The distance travelled by each individual (deltaPCA) depended on the environment of exposure as well, with an effect of origin of the transplant too (Fig. 4). For both factors, the shallow level induced less change (Fig. 4).

Results show differences in the mean responses of species along axes of morphological variation. The random effect structure resulting from model selection included Genotype for all the models but not rack. Species was selected for eight models, for which intra-specific variance across genotypes was comparable or greater than variance due to species identity for most cases (Figure 5 and Table SM1). This suggests that individual plasticity of each genotype varies in comparable magnitude with variation in the response occurred among species, as confirmed by the reaction norm plots (Figure SM2 and SM3).

**Figure 5 –.**
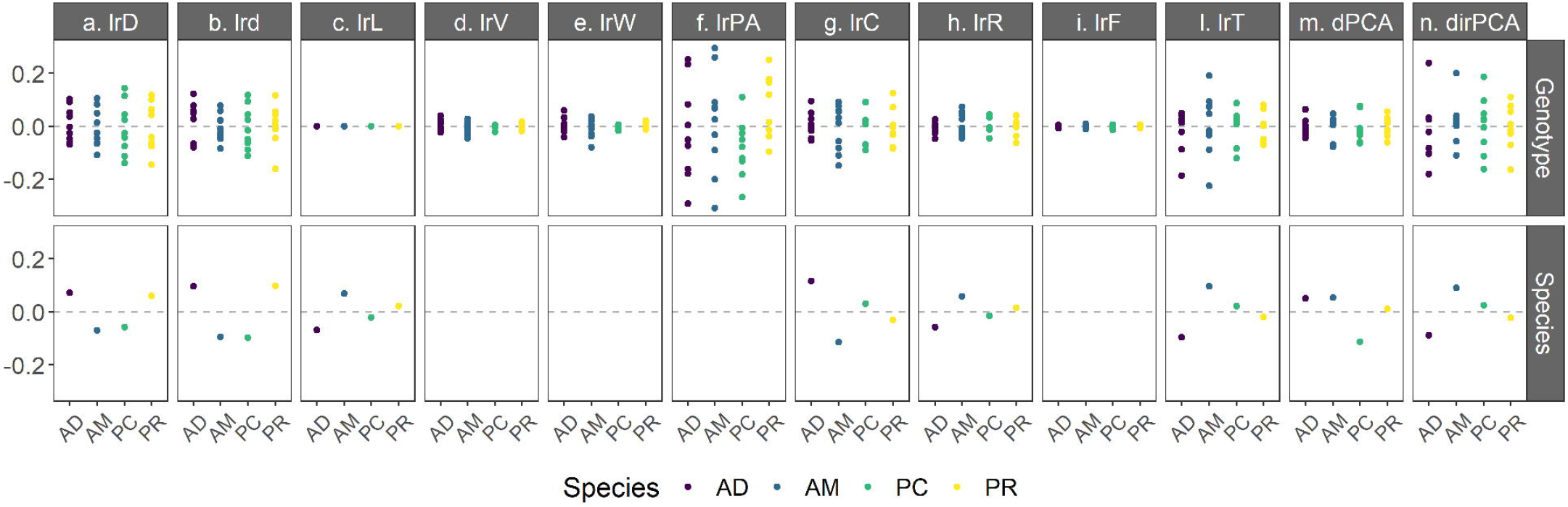
Random effects in selected models. Points correspond to Genotype and Species effects in the models selected. Models without effects of Species do not have this variable in their selected random effect structure. AM = *Acropora muricata*, AD = *Acropora divaricata*, PC = *Porites cylindrica*, PR = *Porites rus*.

Bonferroni corrected significancy of the fixed effects in the full models fitted for each response variables are qualitatively consistent with model selection results (see SM “Full models fitted for each individual trait”, Table SM3 and Figure SM9).

## Discussion

This study showed evidence for phenotypic plasticity as the main driver of variation in coral morphology. Transplant destination consistently induced the greatest difference in shape change, with the highest differences among *Acropora* spp.. The origin of the corals (*i*.*e*., the site where corals lived before transplantation) had minimal effects on some morphological traits, including changes in the multidimensional trait space. We failed to detect a significant interaction between origin and destination, which suggest that local adaptation does not depend on the environment of previous exposure and on any filtering that may have happened there. Similarly to other transplant experiments (12), genotype by environment interactions (intraspecific variation) were detected. Given our small effect of origin and the consistency of the reaction norms, our study suggests that the morphological variability found in coral reefs is more strongly linked to plasticity than environmental filtering.

When comparing environmental conditions of the two experimental sites in the lagoon, at the shallow site mean light conditions were twice as high, water motion was greater (though overall negligible), and daily variability of temperature and light were higher. These conditions are common in shallow reefs and can be stressful for corals during thermal anomalies (37). Such variability in environmental conditions provides a plausible explanation for the lower survival rate, especially among the *Acropora* spp., which are particularly sensitive to high temperature exposure (38) and for which we can see a subtle, even though not significant signal (Fig.2). Apart from survival, in the deep, more stable, environment we observed a higher diversification in morphologies. This was shown by the larger morphospace occupied by corals transplanted to the deep site (Fig.3 and SM7). The shallow site covered a smaller extent of morphologic traits combinations, as showed by the restricted portion of the morphospace occupied by the shallow site at the end of the experiment (Fig.3 and SM7).

Looking at differences between genera, and pooling together survival and morphological traits results, *Porites* spp. and *Acropora* spp. displayed different responses. Variation between species within genus was small, suggesting that plasticity in the traits analysed may be conserved at higher taxonomic groups. *Porites* spp. grew less (except at the base of the skeleton) and had smaller changes in complexity and compactness, but higher survival rate. This direction of change was expected as this genus is prone to develop basally before growing branches (27, 39). *Acropora* spp., on the other hand, had higher changes in traits, especially at the deep site where there was higher survival rate. This could be due to the higher growth rate in *Acropora* spp. (27) and the tendency to develop open branches and top-heavy morphologies (20,22). Although this variation in coral morphologies is visible when colony size is bigger, our results show that different environment can change the rate and direction of change in relatively small time (*i*.*e*., less than 1.5y), which is impressive given the lifespan of corals.

Since corals are autogenic ecosystem engineers, they promote habitat diversification with their diversification of their morphology (2)(40) Our results indicate no trade-off between survival and habitat diversification, but rather that this relationship is additive. The higher the survival, the more traits changed, and the higher the habitat diversification ability for the ecosystem. In contrast, corals transplanted to the shallow site survived less and displayed less variation in traits, which in a reef may result in less varied habitat provision.

Origin had no effect on survival and had a small effect in only three of the morphologic traits considered. The fact that the experiment was carried out in an enclosed lagoon may have facilitated the gene flow between deep and shallow populations, compared to less connected exposed reef crests and slope depths (41). High connectivity may therefore explain the minor effect of origin in this experiment, but the two sites were nonetheless approximately 150m apart. The fact that trait changes were marginally higher in genotypes selected from the deep sites, that there was no effect of the direction of transplant, and that there was no effect on survival might be reassuring for conservation reasons. In fact, a common practice is to transplant coral fragments from other sites in reefs that need restoration and wait for corals to attach and grow (42). If origin and direction of transplantation do not affect coral growth, but the environment does, then one can predict that after some time the same morphological patterns appear in the reef. In the long term though, following corals along generations would be necessary do detect if there might be an adaptive response that will benefit restoration, as other lines of research are currently investigating (43).

The present findings have also important evolutionary implications. The debate surrounding the extended evolutionary synthesis revolves around the role of genetic versus non genetic controls of phenotype (44). Our results suggest stronger environmental control on the morphological phenotype. However, phenotypic plasticity itself can be under genetic control. Interestingly, we found that plasticity was higher among *Acropora* corals, which is the most speciose genus among Scleractinian corals with more than 150 described species (20,45) and more still being discovered. In fact, the ability to sustain morphological diversity through phenotypic plasticity may affect evolutionary rates (46). As niche constructors, corals can affect reef habitat diversity, and consequently selection patterns on themselves and other organisms. Our results suggest that this can happen mainly through phenotypic plasticity. This mechanism could be important for understanding coral and reef evolution as could be related to speciation patterns and niche construction processes (44). Testing whether the patterns found for the four species in this study generalize across other ecosystem engineers will be important.

Structure plays a critical role in the ecology of any ecosystem. In coral reefs in particular, high structural complexity increases fish and invertebrate biomass and biodiversity (47,48). As a result, algal cover decreases possibly due to the increase of herbivores (49). Structural complexity also correlates with coral diversity and abundance (50) and with the reef ability of recovering faster after disturbance (51). Here we show that the shape of structure is highly responsive to shifting environmental conditions. Given the links between structure and function in this ecosystem (47-51), our results highlight how function is highly sensitive to shifting environmental conditions.

This study illustrates the importance of transitioning towards quantitative morphologic traits for detecting variation in shape complexity. This approach opens opportunities to detect continuous differences along gradients, and to gain a deeper insight into the drivers of differences in structure builders of complex habitats. Moreover, a significant benefit of using functional morphologic traits derived from planar area representations is that they can be applied to archived images taken during other transplant experiments, making a generalization of these findings possible. Improved understanding of the relationship between the traits computed here and the 3D morphologic traits described in Zawada et al. (22) could help us move further forward in understanding the ecological consequences of plasticity in coral morphologies (23).

Overall, this study showed that coral ecosystem engineering traits are phenotypically plastic and differ across individuals. The origin of the individuals affected only some of the traits, and marginally. Different local environmental conditions consistently induce different coral morphologies, and consequently varied habitat availability in reef ecosystem. Hence, our results suggest reef zonation and functioning are largely driven by coral phenotypic plasticity. As environmental condition shift across the world, these are likely to be quickly reflected in coral reef structure, with ecological consequences for this already threaten system.

## Ethics

This research was approved by the Ministry of Fisheries and Agriculture of Maldives (protocol numbers: (OTHR)30-D/INDIV/2016/537 and (OTHR)30-D/lNDlV/2018/739.

## Data accessibility

The code and processed data supporting this article will be made public upon peer-reviewed publication in GitHub at https://github.com/VivBramb/ShapingCoralTraits. Access can be granted upon request to the corresponding author.

## Authors’ contribution

VB: Conceptualization; Data curation; Formal analysis; Investigation; Methodology; Project administration; Software; Supervision; Visualization; Writing – original draft; Writing – review & editing. MB: Data curation; Methodology; Resources; Writing – review & editing. ID: Data curation; Resources. JM: Conceptualization; Investigation; Methodology; Writing – review & editing. DM: Data curation; Resources; Writing – review & editing. CP: Data curation; Resources. MD: Conceptualization; Data curation; Funding acquisition; Investigation; Methodology; Project administration; Resources; Supervision; Writing – review & editing. All authors gave final approval for publication and agree to be held accountable for the work performed therein.

## Acknowledgements

We thank Kevin Laland and James Guest for comments and feedback on a previous draft of this manuscript. We thank Mia Hoogenboom for feedback on the design of the experiment. We thank MaRHE center for support in the field and the Behaviour and Biodiversity group at University of St Andrews for feedback and suggestions during the data analysis.

## References

1. West-Eberhard MJ. Developmental plasticity and evolution. 2003.

2. Jones CG, Lawton JH, Shachak M. Organisms as ecosystem engineers. Oikos. 1994;69(3):373–86.

3. Odling-Smee FJ, Laland KN, Feldman MW. Niche Construction. Am Nat. 1996 Apr 1;147(4):641–8.

4. Callaway RM, Walker LR. Competition and facilitation: a synthetic approach to interactions in plant communities. Ecology. 1997;(78):1958–65.

5. Rovira AD. Interactions Between Plant Roots and soil microorganisms. Annu Rev Microbiol. 1964;19(1):241–66.

6. Miner BG, Sultan SE, Morgan SG, Padilla DK, Relyea RA. Ecological consequences of phenotypic plasticity. Trends Ecol Evol. 2005;20(12):685–92.

7. Donohue K. Why ontogeny matters during adaptation: Developmental niche construction and pleiotorpy across the life cycle in arabidopsis thaliana. Evolution (N Y). 2014;68(1):32–47.

8. Keddy PA. Assembly and response rules:two goals for predictive community ecology. J Veg Sci. 1992;3:157–64.

9. Bradshaw AD. Environment and phenotypic plasticity. Brookhaven Symp Biol. 1974;25(75–94).

10. Gattuso J-P, Allemand D, Frankignoulle M. Photosynthesis and Calcification at a Cellular, Organismal and Community Levels in Coral Reefs: A Review of Interactions and Control by Carbonate Chemistry. Am Zool. 1999;39:160–83.

11. Hoogenboom MO, Connolly SR. Defining fundamental niche dimensions of corals:synergistic effects of colony size, light, and flow. Ecology. 2009;90(3):767–80

12. Todd PA. Morphological plasticity in scleractinian corals. Biol Rev. 2008;83(3):315–37.

13. Dustan P. Growth and form in the reef-building coral Montastrea annularis. Mar Biol. 1975;33:101–7.

14. Jaubert J. Light, metabolism and growth forms of the hermatypic scleractinian coral Synaraea convexa Verrill in the lagoon of Moorea (French Polynesia). Proc Third Int Coral Reef Symp. 1977;1:483–488.

15. Ow YX, Todd PA. Light-induced morphological plasticity in the scleractinian coral Goniastrea pectinata and its functional significance. Coral Reefs. 2010;29(3):797–808.

16. Stambler N, Dubinsky Z. Corals as light collectors: an integrating sphere approach. Coral Reefs. 2005;24(1):1–9.

17. Hoogenboom MO, Connolly SR, Anthony KRN. Interactions between morphological and physiological plasticity optimize energy acquisition in corals. Ecology. 2008;89(4):1144–54.

18. Chappell J. Coral morphology, diversity and reef growth. Nature. 1980 Jul;286(5770):249–52.

19. Sebens KP, Witting J, Helmuth B. Effects of water flow and branch spacing on particle capture by the reef coral Madracis mirabilis (Duchassaing and Michelotti). J Exp Mar Bio Ecol. 1997 Apr;211(1):1–28.

20. Veron JEN, Stafford-Smith M. Corals of the world. Townsville,Australia: Australian Institute of Marine Sciences; 2000. 1350 p.

21. Wallace C. Staghorn Corals of the World. CSIRO Publishing; 1999.

22. Zawada KJA, Dornelas M, Madin JS. Quantifying coral morphology. Coral Reefs. 2019;38:1281–1292.

23. Zawada KJA, Madin JS, Baird AH, Bridge TCL, Dornelas M. Morphological traits can track coral reef responses to the Anthropocene. Funct Ecol. 2019;33:962–75.

24. Todd PA, Sidle RC, Lewin-Koh NJI. An aquarium experiment for identifying the physical factors inducing morphological change in two massive scleractinian corals. J Exp Mar Bio Ecol. 2004;299(1):97–113.

25. Bruno JF, Edmunds PJ. Clonal Variation for Phenotypic Plasticity in the Coral Madracis Mirabilis. Ecology. 1997;78(7):2177–90.

26. Bruno JF, Edmunds PJ. Metabolic consequences of phenotypic plasticity in the coral Madracis mirabilis (Duchassaing and Michelotti): The effect of morphology and water flow on aggregate respiration. J Exp Mar Bio Ecol. 1998;229(2):187–95.

27. Madin JS, Anderson KD, Andreasen MH, Bridge TCL, Cairns SD, Connolly SR, et al. The Coral Trait Database, a curated database of trait information for coral species from the global oceans. Sci Data. 2016;3:160017.

28. Smith LW, Wirshing HH, Baker AC, Birkeland C. Environmental versus Genetic Influences on Growth Rates of the Corals Pocillopora eydouxi and Porites lobata (Anthozoa: Scleractinia). Pacific Sci. 2008;62(1):57–69.

29. Jokiel P. Coral growth:buoyant weight. Coral Reefs. 1978529–541.

30. Fulton CJ, Bellwood DR. Wave-induced water motion and the functional implications for coral reef fish assemblages. Limnol Oceanogr. 2005;50(1):255–64.

31. Rasband W. ImageJ. Bethesda, Maryland, USA: U.S. National Institutes of Health; 2014.

32. Borchers HW. pracma: Practical numerical math functions. R package; 2019. p. 1–393.

33. Sevçíková H, Percival DB. fractaldim: Estimation of fractal dimensions. R-Package. 2015.

34. Zuur A, Ieno EN, Walker N, Saveliev AA, Smith GM. Mixed effects models and extensions in ecology with R. Springer Science & Business Media; 2009.

35. R Core Team. R: A language and environment for statistical computing. 2018.

36. Bates D, Mächler M, Bolker BM, Walker SC. Fitting Linear Mixed-Effects Models Using lme4. 2015;67(1).

37. Glynn PW. Coral reef bleaching: Facts, hypotheses and implications. Glob Chang Biol. 1996;2(6):495–509.

38. Hoogenboom MO, Frank GE, Chase TJ, Jurriaans S, Álvarez-Noriega M, Peterson K, et al. Environmental Drivers of Variation in Bleaching Severity of Acropora Species during an Extreme Thermal Anomaly. Front Mar Sci. 2017 Nov 27;4.

39. Yap HT, Molina RA. Comparison of coral growth and survival under enclosed, semi-natural conditions and in the field. Mar Pollut Bull. 2003;46(7):858–64.

40. Jones CG, Lawton JH, Shachak M. Organisms as Ecosystem Engineers. Oikos. 1994;69(3):373–86.

41. Benzie JAH, Haskell A, Lehman H. Variation in the genetic composition of coral (Pocillopora damicornis and Acropora palifera) populations from different reef habitats. Mar Biol. 1995;121:731–9.

42. Barton JA, Willis BL, Hutson KS. Coral propagation: a review of techniques for ornamental trade and reef restoration. Rev Aquac. 2017;9(3):238–56.

43. van Oppen MJH, Oliver JK, Putnam HM, Gates RD. Building coral reef resilience through assisted evolution. Proc Natl Acad Sci. 2015;112(8):1–7.

44. Laland KN, Uller T, Feldman MW, Sterelny K, Muller GB, Moczek A, et al. The extended evolutionary synthesis: its structure, assumptions and predictions. Proc R Soc B Biol Sci. 2015;282(1813):20151019.

45. Wallace CC, Willis BL. Systematics of the coral genus Acropora: Implications of new biological findings for species concepts. Annu Rev Ecol Syst. 1994;25(March 2015):237–62.

46. Matthews B, De Meester L, Clive GJ, Ibelings BW, Bouma TJ, Nuutnen V, et al. Under niche construction: an operational bridge between ecology, evolution, and ecosystem science. Ecol Monogr. 2014;84(2):245–63.

47. Darling ES, Graham NAJ, Januchowski-Hartley FA, Nash KL, Pratchett MS, Wilson SK. Relationships between structural complexity, coral traits, and reef fish assemblages. Coral Reefs. 2017;36(2):561–75.

48. Stella JS, Pratchett MS, Hutchings PA, Jones GP. Coral-associated invertebrates: diversity, ecological importance and vulnerability to disturbance. Oceanogr Mar Biol An Annu Rev. 2011;49:43–104.

49. Graham NAJ, Nash KL. The importance of structural complexity in coral reef ecosystems. Coral Reefs. 2013;32(2):315–26.

50. Torres-Pulliza D, Dornelas MA, Pizarro O, Bewley M, Blowes SA, Boutros N, et al. A geometric basis for surface habitat complexity and biodiversity. Nat Ecol Evol. 2020;

51. Dajka J-C, Wilson SK, Robinson JPW, Chong-Seng KM, Harris A, J.Graham NA. Uncovering drivers of juvenile coral density following mass bleaching. Coral Reefs. 2019;38(4):637–49.

