## Supplemental material for "Shaping coral traits: plasticity more than filtering"

Supplementary material for *Shaping coral traits*

Figure SM4 – Pair plot for the trait analysed.	8

[Figure SM8 – Genus and Destination interaction plots for significant interactions	12](https://d.docs.live.net/3c63ca7f770a002d/RTE%20manuscript/RTE_analysis_SM.docx#_Toc74823165)

### Reaction norms and raw data

Figure SM1 (next page) – Reaction norms in reciprocal transplant experiments. These plots represent how reaction norms are usually represented and offer keys of interpretations specifically for reciprocal transplant experiments (when the site of origin of the genotype can also influence the norm). In the first panel, figure 1a and 1b represent the reaction of one individual genotype. 1a shows a genotype that display the same trait value among sites (canalized norm), while 1b represent a plastic trait, which display different values according to the environment. In the second panel, 2 genotypes per plot are present. In the second panel, different genotypic effects are displayed. The genotypes of figure 2a) and 2b) are influenced in the same fashion by the environment (i.e. the difference in the trait between sites is the same), but while in 1a) the norms are identical, in 2b) they are parallel. In figure 2c) and 2d) there are genotype by environment interactions. In 2c) the norms of the genotypes differ for magnitude, while in 2d) they differ in direction of response. In the third panel, effects of the origin site, represented by different line type, are introduced. Norms in 3a) show an effect of the interaction between site of origin and environment, since norms of individuals with the same origin have the same reaction norms when exposed to the same environments. Norms in 3b) show when differences in the magnitude are not affected by origin. Norms in 3c) show the null effect of origin on the direction of the norms. In the fourth panel, taxonomic groups are included as different colours. In 4a) norms show an effect of origin, rather than taxonomic groups. In 4b) there is an effect of the taxonomic group, rather than origin. In 4c) there is an interaction between origin, taxonomic group and environment.

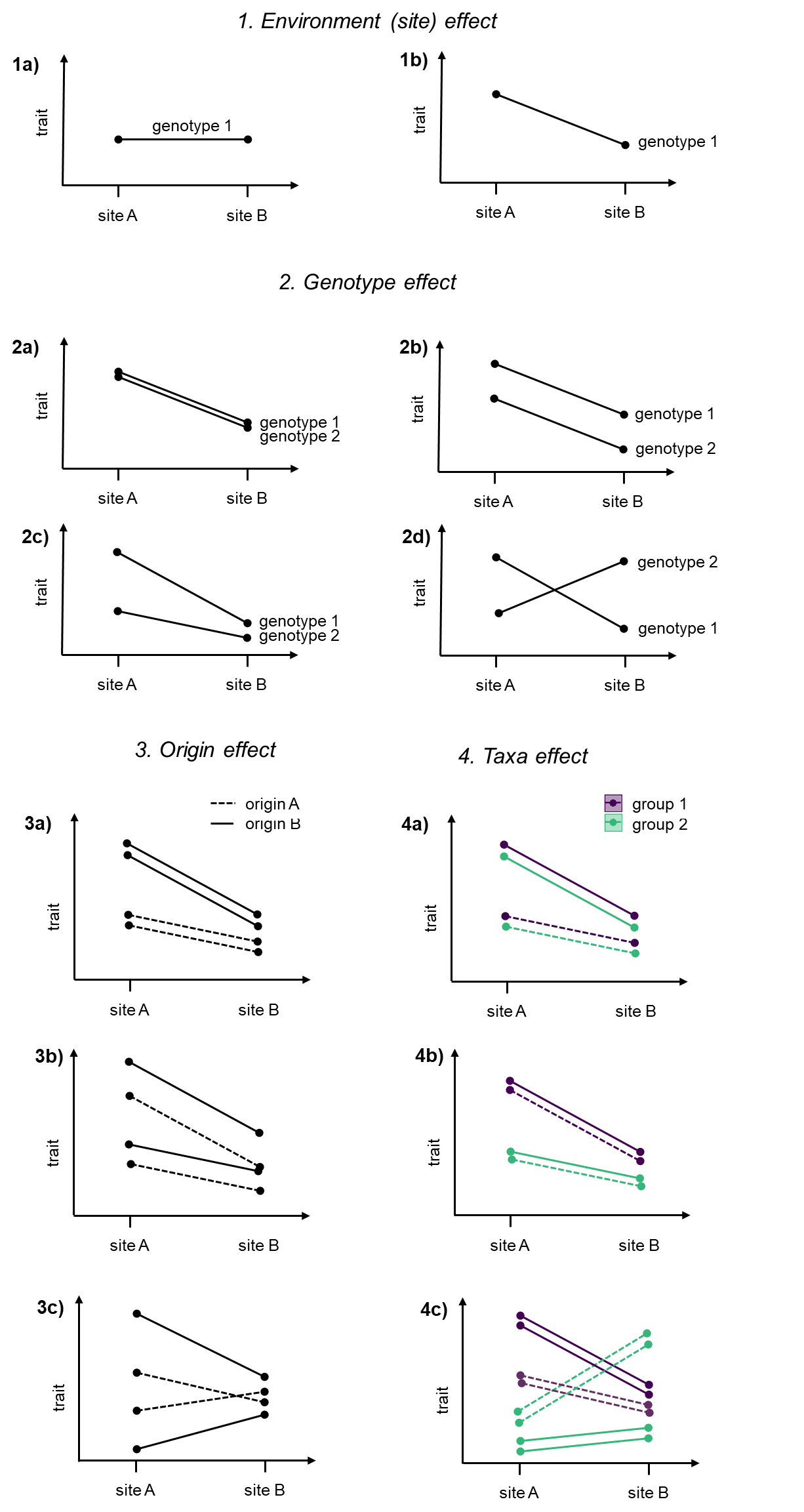

**Figure SM2 Species trait distributions at the beginning and at the end of the experiment for the traits measured in situ (n=218).** Values of each trait are represented color-coded by species at the beginning (t0) and at the end (t1) of the experiment. Both values (dots) and distributions (violins) are shown. The coloured lines link the means of the trait values of each species before and after the experiment under each experimental treatment. For every trait, data is divided by site of Origin of the source colony (deep and shallow Origin, as different columns in each panel) and site of Destination (deep and shallow Destination, as different rows). If slopes are consistent among trait panels, then the nubbins did not display morphological plasticity, nor showed an effect of the Origin on the genotype of the colonies. If patterns of the lines change among columns, then there is a strong determination of the morphs based on the Origin of the colony. If patterns of the lines change among rows, then the environment affected the trait values that changed due to morphological plasticity. For example, in the graph for maximum basal diameter (a), while the 2 columns have almost the same lines pattern (i.e. the response did not depend on the Origin nor the Destination), the two *Porites spp.* (PC and PR) show different pattens when comparing between rows and among columns (i.e. the Destination seems to affect this trait).

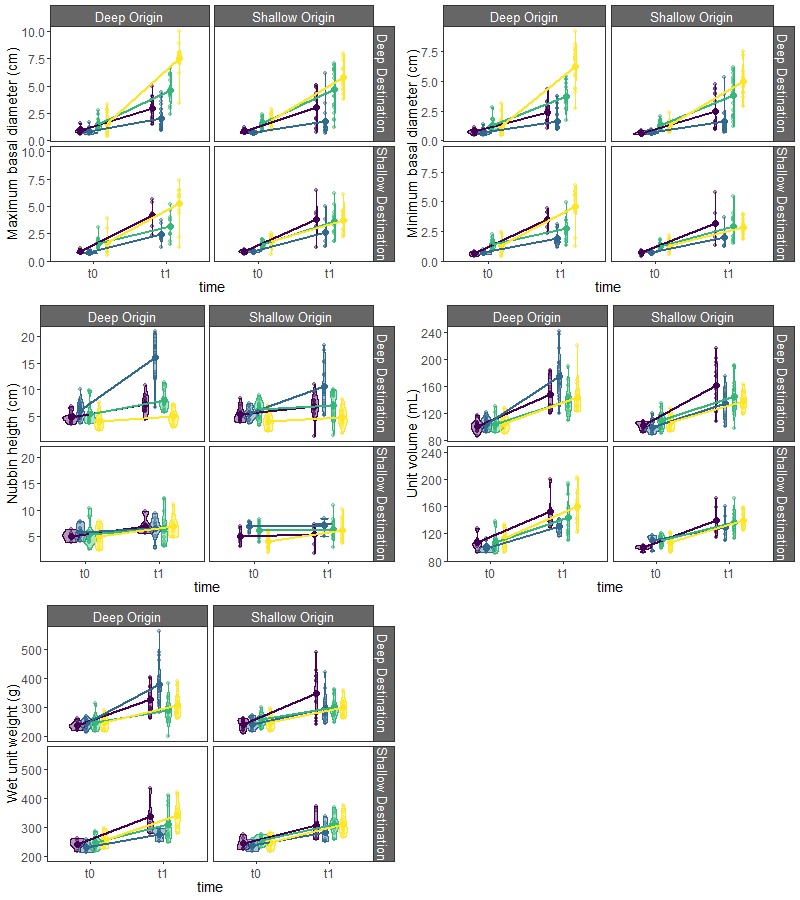

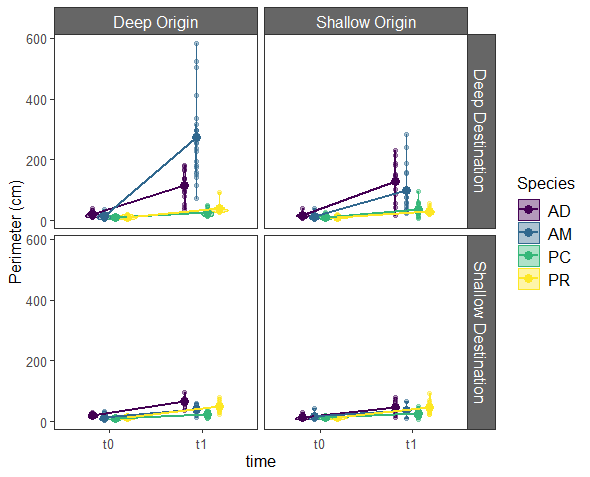

**b)**

**c)**

**a)**

**d)**

**e)**

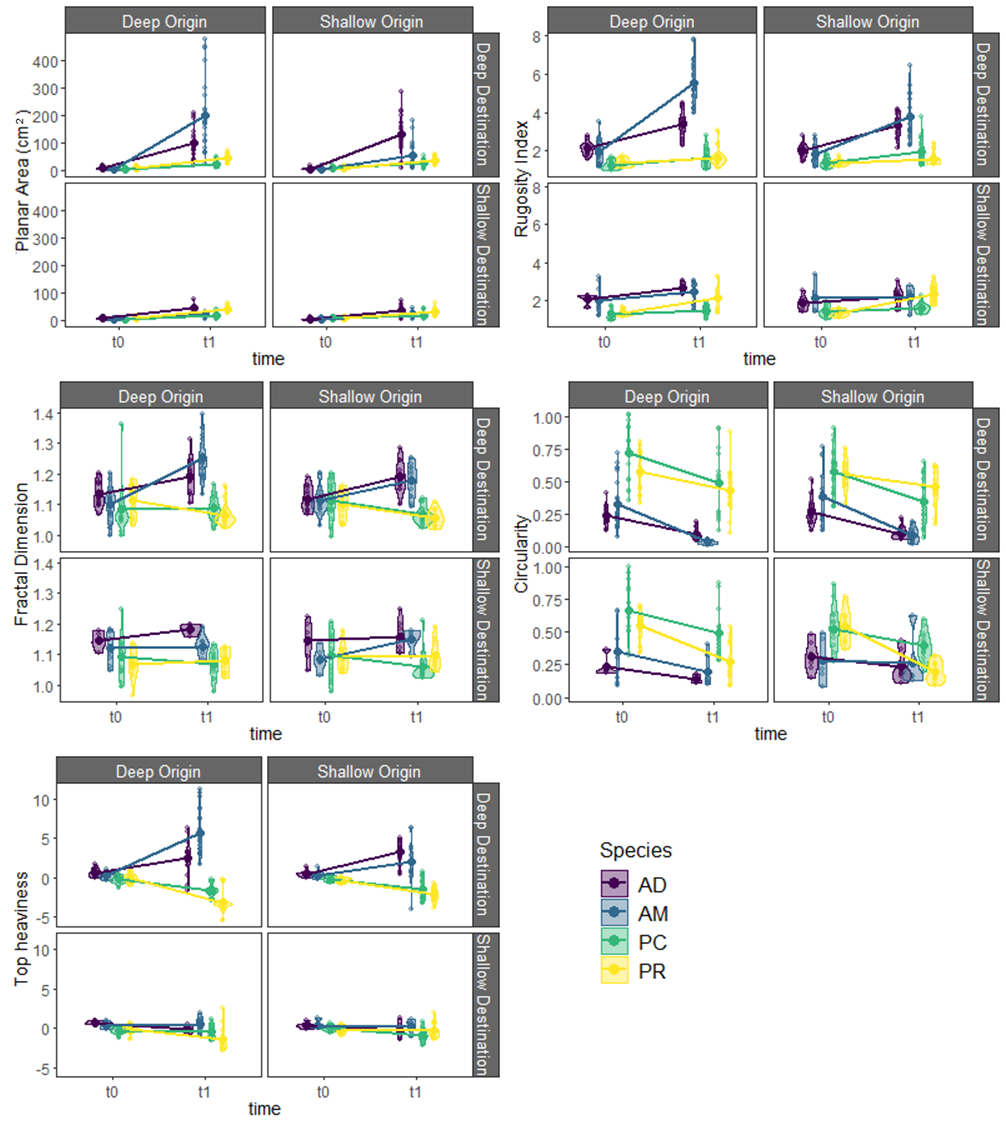

Figure SM2 (continuation) - Species trait distributions at the beginning and at the end of the experiment for the traits derived from the nubbin outline (n = 217). Values of each trait are represented color-coded by species at the beginning (t0) and at the end (t1) of the experiment. Both values (dots) and distributions (violins) are shown. The coloured lines link the means of the trait values of each species before and after the experiment under each experimental treatment. For every trait, data is divided by site of Origin of the source colony (deep and shallow Origin, as different columns in each panel) and site of Destination (deep and shallow Destination, as different rows). For interpretation, see previous page.

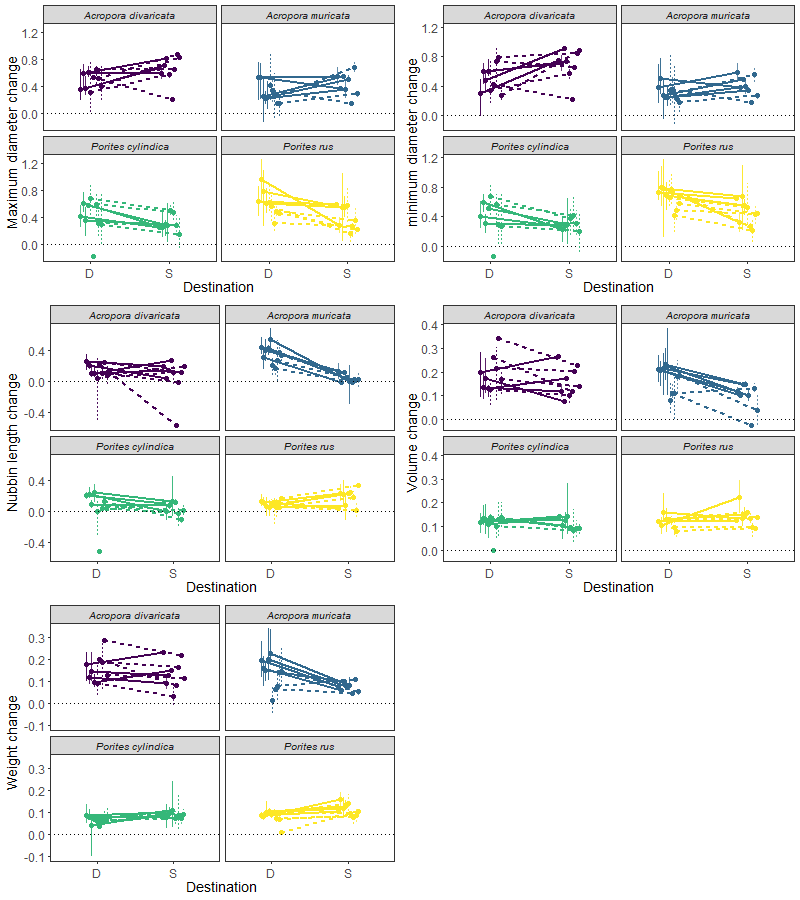

Figure SM3 – Species reaction norms for the change in traits (log ratio) measured in situ (for interpretation, refer to Figure SM1). Vertical lines represent the range of the trait values of each genotype at each Destination. Dots represent mean value of traits per each genotype at each Destination site. Mean values corresponding to the same genotype are connected by a line. Line type changes according to Origin site (dotted for shallow Origin and dotted for deep Origin). Data is color-coded by species.

Figure SM3 – Species reaction norms for the change in traits (log ratio) derived from nubbin outlines (for interpretation, refer to Figure C.1). Vertical lines represent the range of the trait values of each genotype at each Destination. Dots represent mean value of traits per each genotype at each Destination site. Mean values corresponding to the same genotype are connected by a line. Line type changes according to Origin site (dotted for shallow Origin and dotted for deep Origin). Data is color-coded by species.

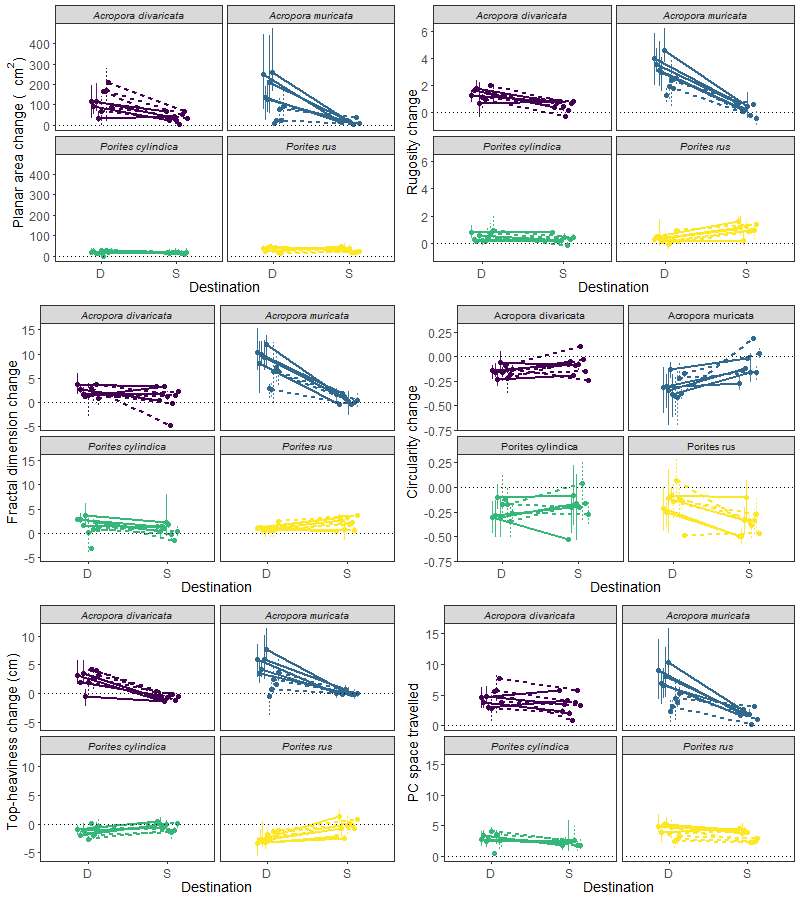

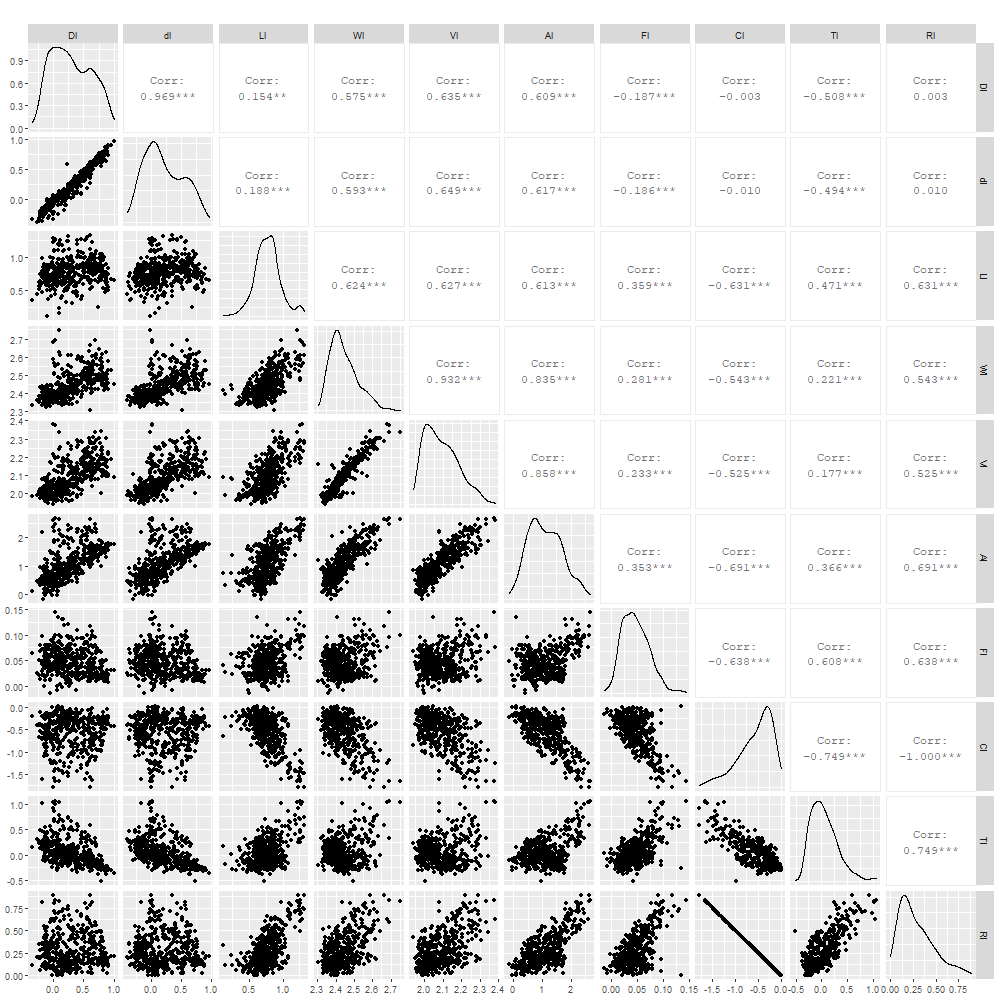

Figure SM4 – Pair plot of the trait analysed. Variables have been log transformed. D = maximum diameter (cm), d = minimum diameter (cm), L = length (cm), V = volume (mL), W = weight (g), A = planar area (cm2), C = circularity index, F = fractal dimension, R = rugosity index, T = top heaviness.

### PCA results

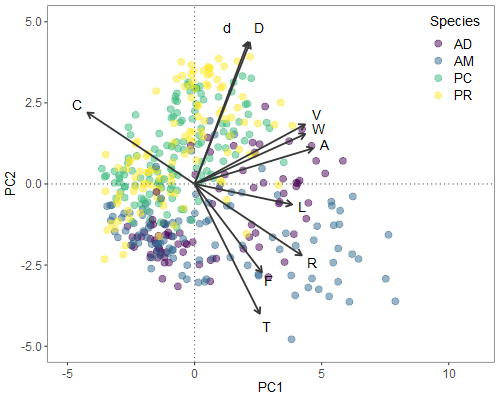

**Figure SM5 – Biplot of the morphospace.** The multidimensional space was built with all the observed trait combinations in the nubbins at the beginning and at the end of the experiment. D = maximum diameter (cm), d = minimum diameter (cm), L = length (cm), V = volume (mL), W = weight (g), A = planar area (cm2), C = circularity index, F = fractal dimension, R = rugosity index, T = top heaviness. Dots are color-coded by species (AD = Acropora divaricata, AM = Acropora muricata, PC = Porites cylindrica, PR = Porites rus). Arrow lengths show how much each trait influence the two principal components of the morphospace (scaled by a factor of 5).

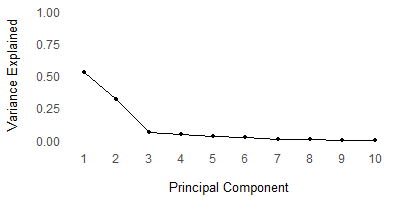

Figure SM6 - Variance explained by each principal component in the morphospace. PC1 explained 53.02% of the variance, PC2 explained 31.43% of the variance. From PC3 less than 6% of the variance was explained.

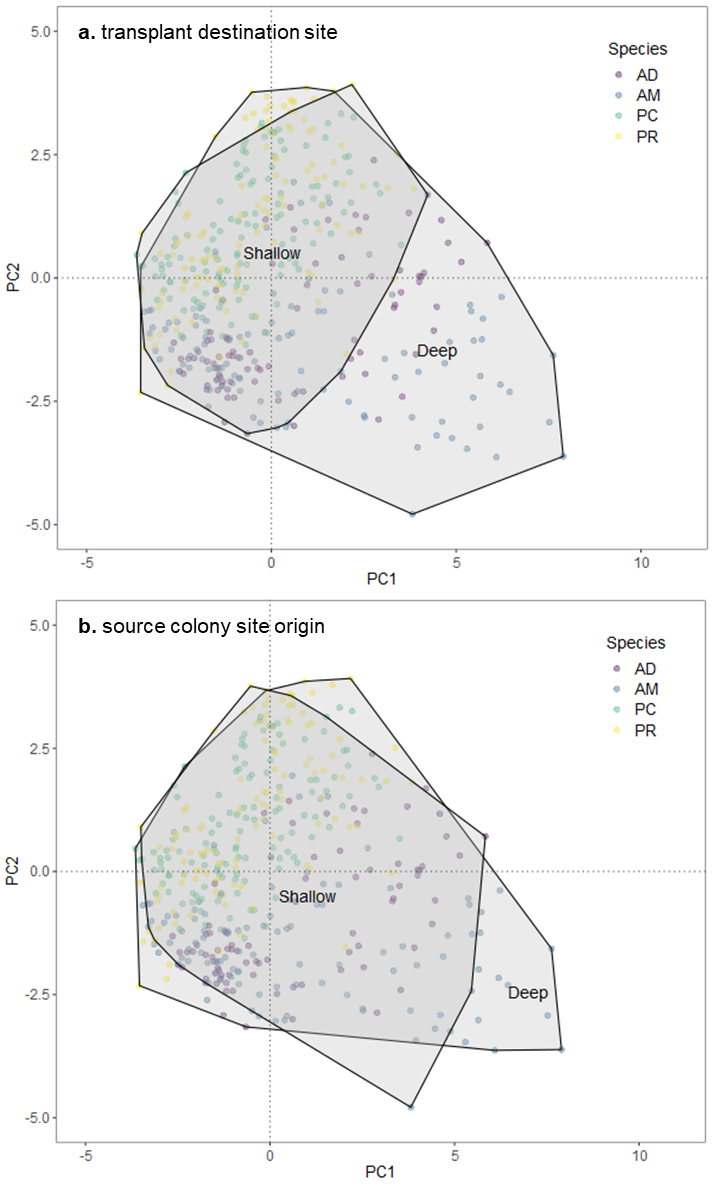

Figure SM7 – Morphological traits space occupied by different site of Origin and Destination. Dots are color-coded by species (*AD = Acropora divaricata, AM = Acropora muricata, PC = Porites cylindrica, PR = Porites rus*). In a) the 2 polygons represent the space occupied by the nubbins that were transplanted on the shallow or the deep site. In b) the 2 polygons represent the space occupied by the nubbins that were originally from the shallow and the deep site.

### Models selected for each individual trait

Maximum and minimum basal diameter models showed that the diameters increased slightly less at the deep site in *Acropora spp.*, but consistently more for *Porites spp.,* regardless of Origin (Figure SM8a-b). Nubbin length and planar area differences across sites were more prominent in Acropora spp., which had higher increases in the deep site (Figure SM8c and f). They were also affected by the coral Origin: genotypes from the deep site population expanded more across the two genera, than genotypes from the shallow site (Figure 4). Volume and weight differences between Destinations were particularly visible in the *Acropora* genus. Nubbins of *Acropora* transplanted into the deep site showed greater increase in the above traits (Figure SM8d-e). Compactness decreased more at the deep site in *Acropora spp.,* regardless of Origin (Figure SM8g). Differences in rugosity and top-heaviness qualitatively mirrored the nubbin volume and weight patterns (Figure 8h and i). While differences between genera were smaller within the shallow site, they were greater in the deep site, where *Acropora spp*. had much higher log ratio responses than *Porites spp*. Model selection for fractal dimension showed that this variable is strictly dependent on Genus with changes *Acropora spp*. being greater regardless of destination and origin (Figure 4).

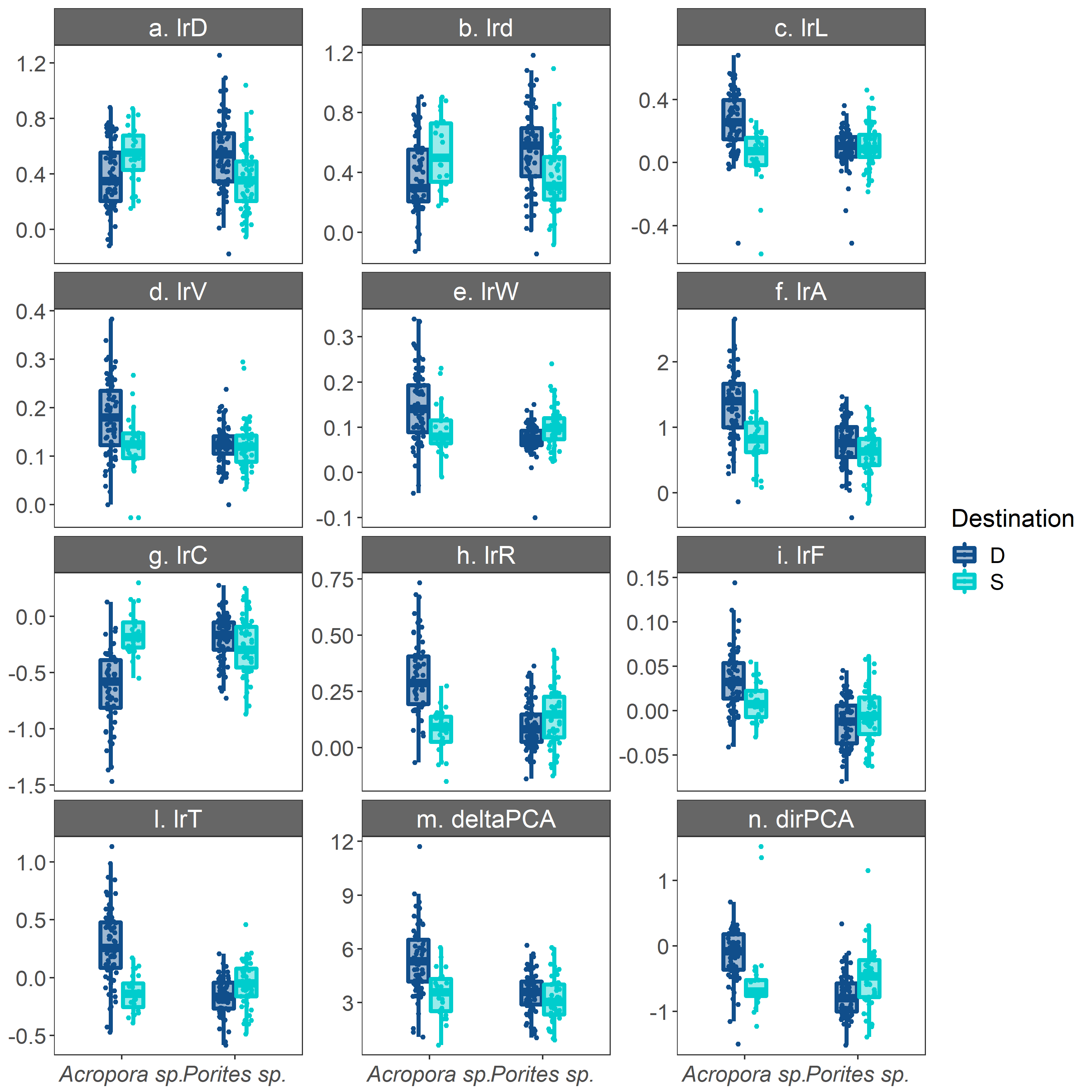

**Figure SM8 – Genus and Destination interaction plots for significant interactions**. Boxplots of trait distributions across genera and transplant destination, with raw data (dots) superimposed. Darker blue represent data transplanted to the Deep site, while lighter blue represent data transplanted to the shallow site. D = maximum diameter (cm), d = minimum diameter (cm), L = length (cm), V = volume (mL), W = weight (g), A = planar area (cm2), C = circularity index, F = fractal dimension, R = rugosity index, T = top heaviness, deltaPCA = distance travelled in the morphospace, dirPCA = direction travelled in the morphospace. Trait change is in log-ratio (‘lr’).

Table SM1 Model selection table. Aikake Information Criterion (AIC) for each fitted model. In bold, the lower AIC for each model fitted.

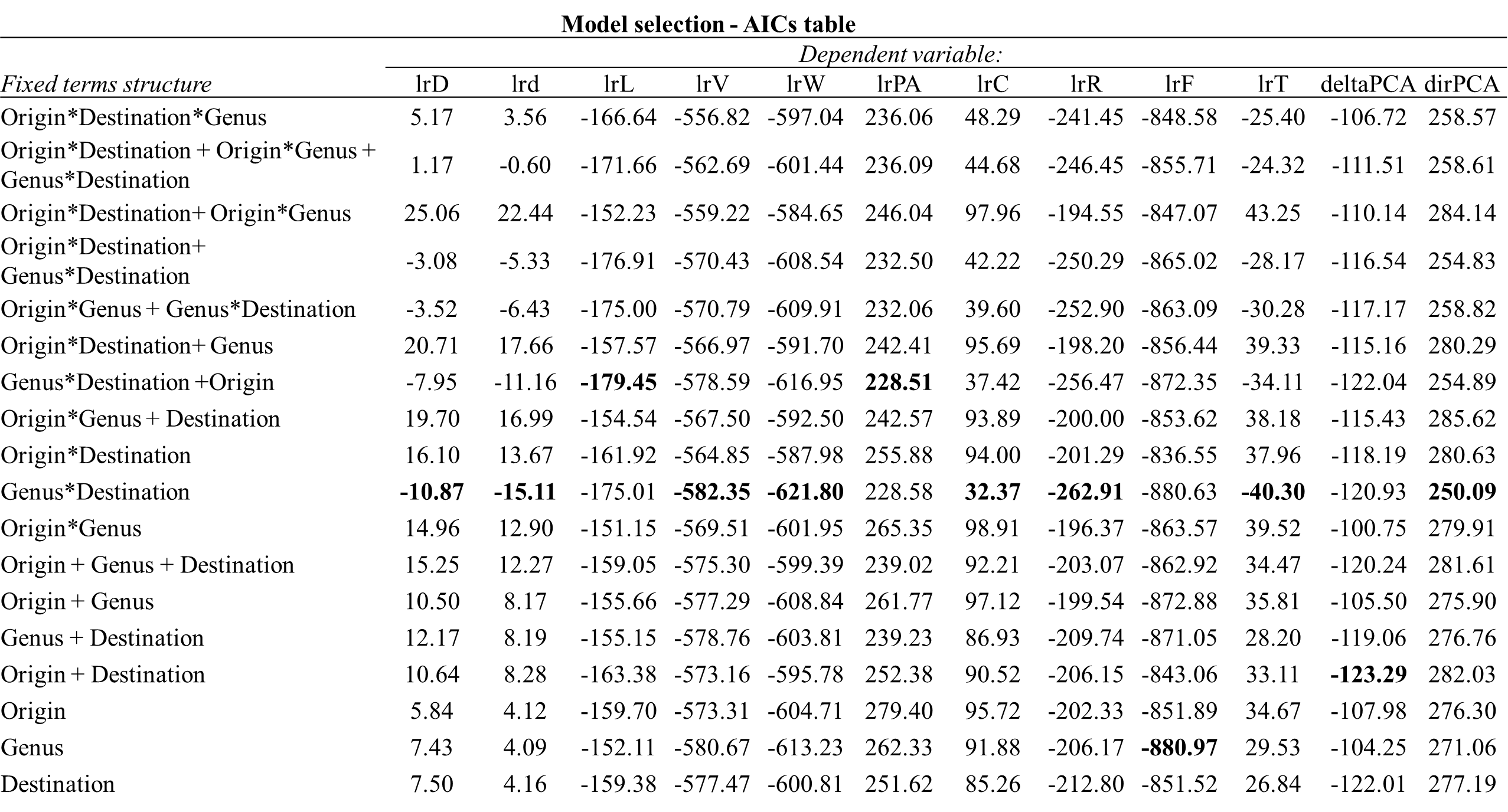

Table SM2 Coefficient estimates (and p.values) for the best models for each of the response variables**.**

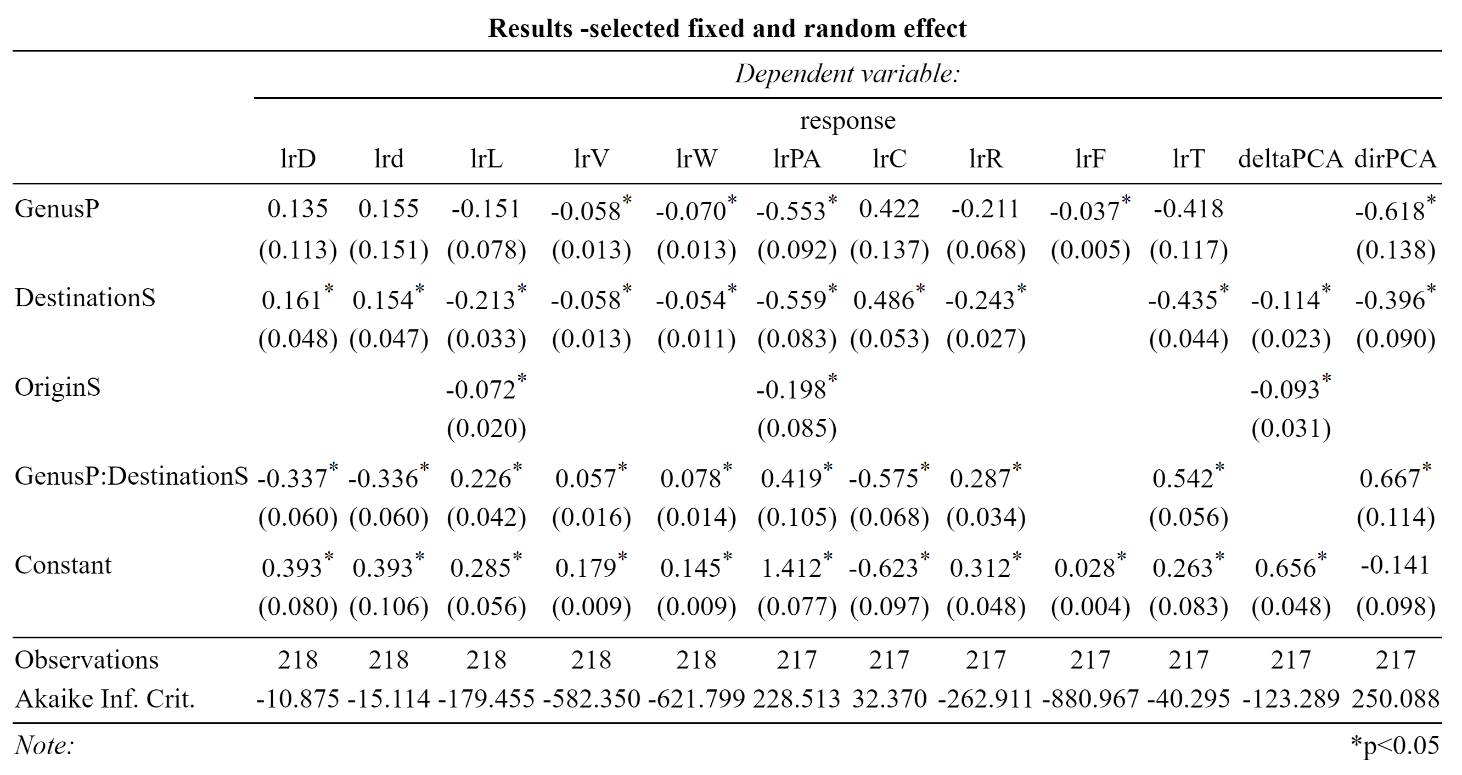

### Full models fitted for the individual traits

Full model results for traits qualitatively confirmed finding from the selected models. In the full models for morphospace analysis, the distance of change (delta PCA) was affected only by Destination. Although there was no effect of Origin, this result confirmed that Destination plays a bigger role than Origin, as effects coefficients from the selected models hinted. The direction of change (dirPCA) in the full model is also affected by a positive interaction of Origin and Destination. Nonetheless, the effects of this interaction are smaller than the other 2 detected.

Table SM3 - Coefficient estimates (and confidence intervals) for full models. Significance levels (*) are Bonferroni-adjusted (n = 8) to account for multiple comparisons.

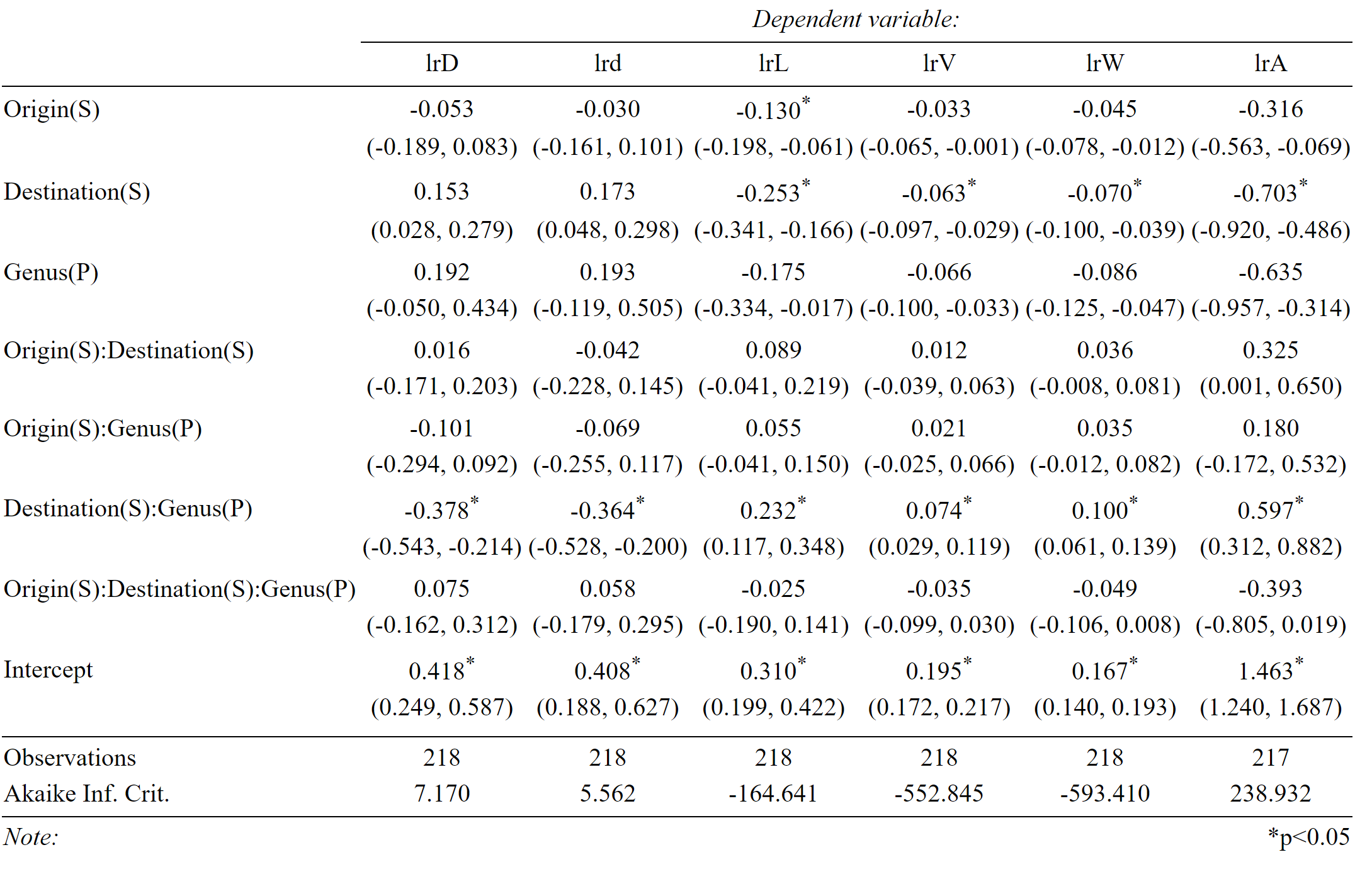

Table SM3 (continuation)- Coefficient estimates (and confidence intervals) for full models. Significance levels (*) are Bonferroni-adjusted (n = 8) to account for multiple comparisons.

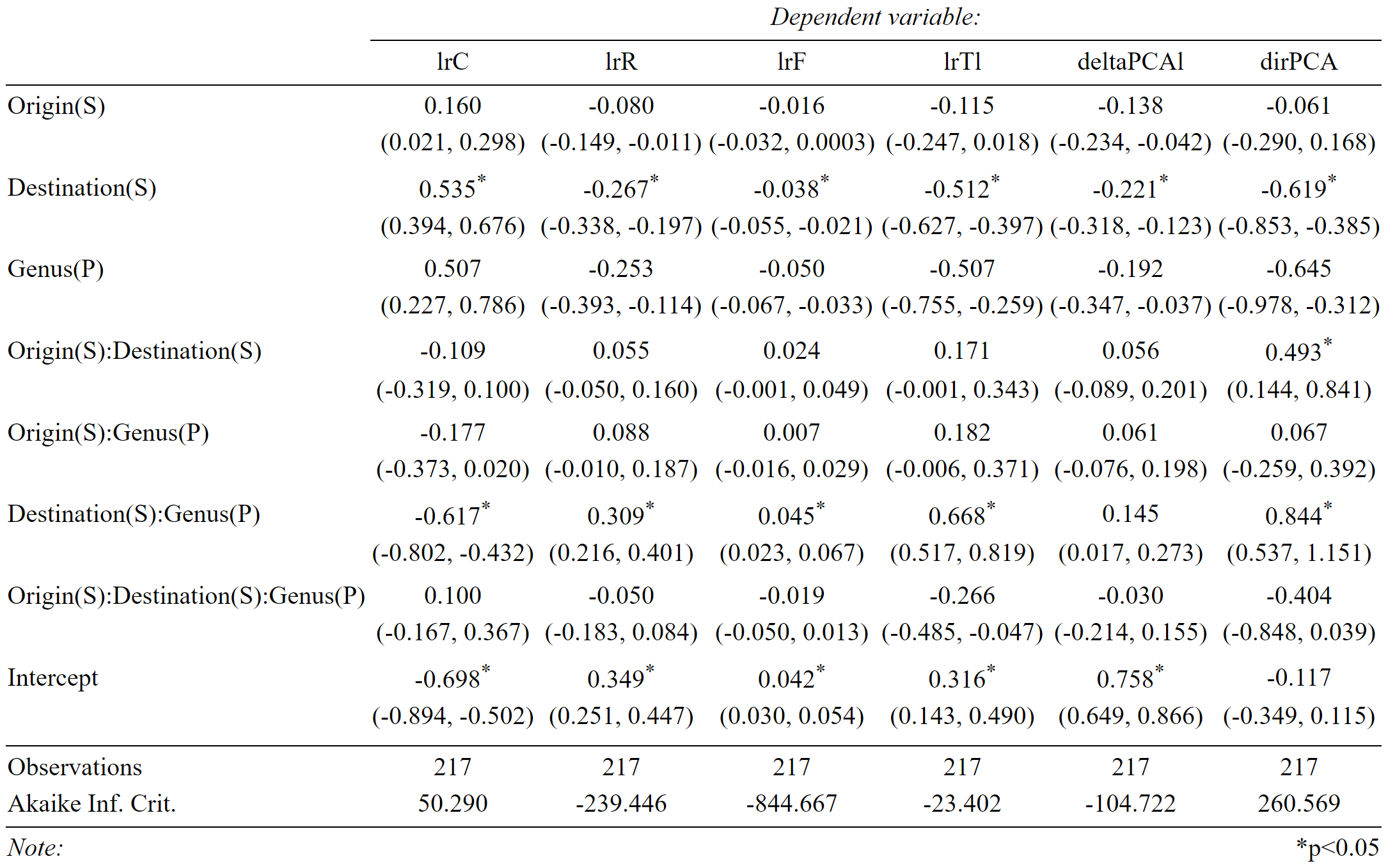

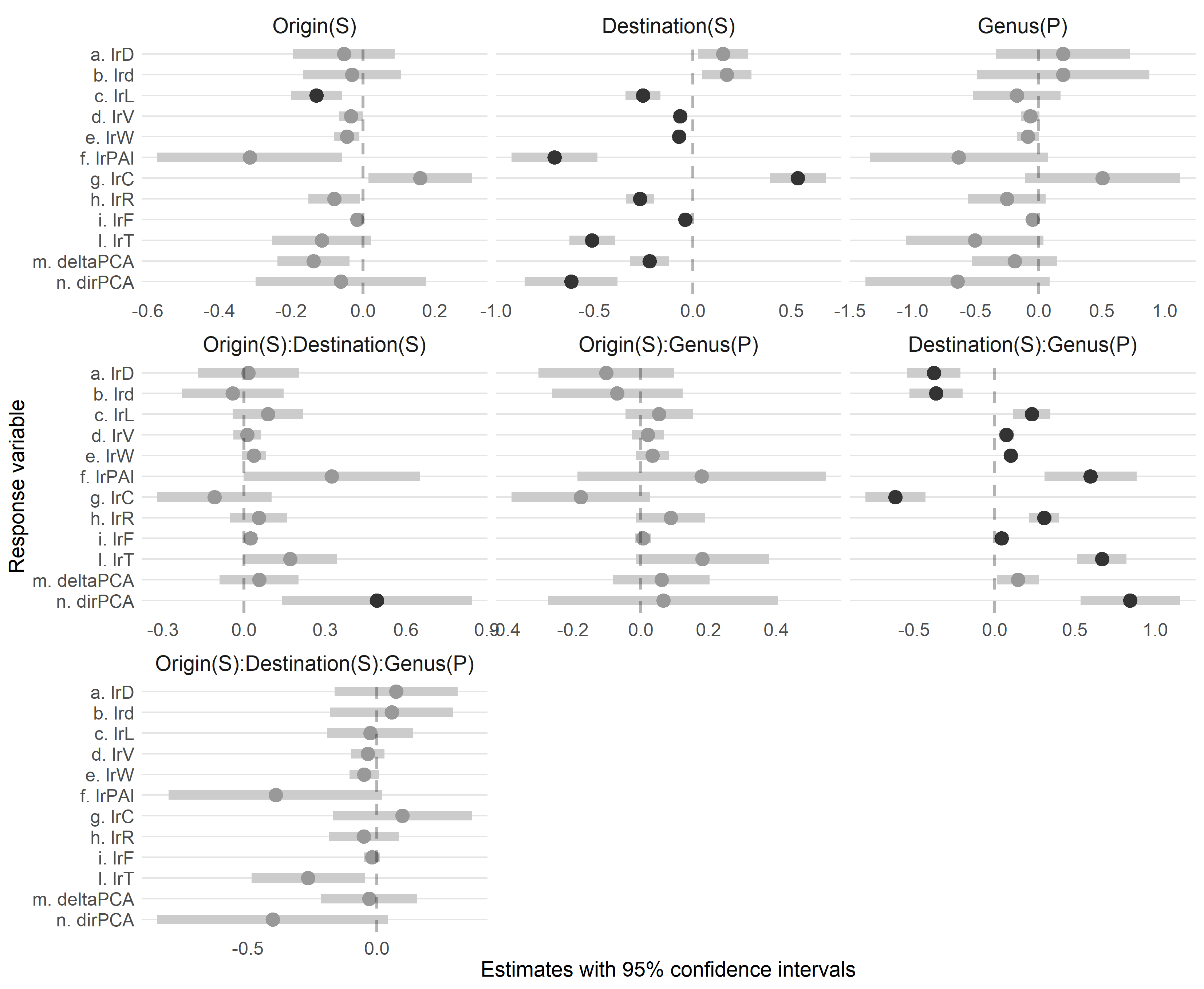

**Figure SM9 - Effect size estimates for fixed factors and interaction terms included in the full models.** Dots represent the estimated effect size of each fixed term in each model and grey bars represent the 95% confidence intervals. Black dots are significant values and significance levels are Bonferroni-adjusted to account for multiple comparisons (n = 8).
